# Rational Design of Self-assembling Artificial Proteins Utilizing a Micelle-Assisted Protein Labeling Technology (*MAPLabTech*): Testing the Scope

**DOI:** 10.1101/2021.08.01.454616

**Authors:** Mullapudi Mohan Reddy, Pavankumar Bhandari, Britto S Sandanaraj

**Affiliations:** Department of Chemistry, Indian Institute of Science Education and Research – Pune; Department of Biology,Indian Institute of Science Education and Research – Pune

## Abstract

Self-assembling artificial proteins (SAPs) have gained enormous interest in recent years due to their applications in different fields. Synthesis of well-defined monodisperse SAPs is accomplished predominantly through genetic methods. However, the last decade witnessed the use of few chemical technologies for that purpose. In particular, micelle-assisted protein labeling technology (MAPLabTech) has made huge progress in this area. The first generation MAPLabTech focused on site-specific labeling of the active-site residue of serine proteases to make SAPs. Further, this methodology was exploited for labeling of *N*-terminal residue of a globular protein to make functional SAPs. In this study, we describe the synthesis of novel SAPs by developing a chemical method for site-specific labeling of a surface-exposed cysteine residue of globular proteins. In addition, we disclose the synthesis of redox- and pH-sensitive SAPs and their systematic self-assembly and dis-assembly studies using complementary biophysical studies. Altogether these studies further expand the scope of MAPLabTech in different fields such as vaccine design, targeted drug delivery, diagnostic imaging, biomaterials, and tissue engineering.

## Introduction

Self-assembling artificial proteins (SAPs) are an interesting class of biomacromolecules that would spontaneously self-associate to forms diverse protein nanostructures^1–5^. These proteins found applications in various fields such as vaccine design^6,7^, targeted drug delivery^8^, in vivo imaging,^9,10^ and tissue engineering.^11,12^ Among the various technologies available to design SAPs; computational protein design^13–16^ has made tremendous progress in the last decade. Similarly, rational protein design^17–19^ and directed evolution technology^20^ also yielded very impressive results. All three technologies utilize a microbial host system to synthesize the target proteins and are therefore restricted to naturally occurring 20 amino acids.

Ashutosh and coworkers introduced a new strategy to synthesize SAPs in which they combined a genetic method with a post-translational modification strategy to synthesize hybrid protein-lipid conjugate^21^. They have shown that these custom-designed proteins self-assemble to form interesting protein nanostructures. Although this method provides opportunities to go beyond the standard 20 amino acids, it is still restricted to a small number of building blocks. In addition, incorporation of every new building block requires elaborate engineering of the host system which restricts the broader utility of this method for various applications. Therefore, it is important to develop a general methodology that can incorporate a wide range of chemical entities onto the self-assembling proteins without compromising salient features of natural proteins such as a single chemical entity and the presence of well-defined functional groups in the 3d-space. Toward that goal, our group invented a new method called ‘Micelle-Assisted Protein Labeling Technology (MAPLabTech)^22–26^. During the last several years, we have utilized MAPLabTech for the design of different families of well-defined monodisperse globular SAPs which include protein amphiphiles^22^, protein-dendron conjugates^23^, protein-peptides conjugates^24^, photo-responsive protein amphiphiles^22^, multi-responsive protein-dendron conjugates^25^, and redox-sensitive protein conjugates^26^. Although extremely powerful, the above method was only applicable to proteins that belong to the serine protease family^22–26^. This is a serious limitation and therefore restricts the use of this method for a variety of applications such as vaccine design^6,7^ and antibody-drug conjugates.^27^ To improve the scope of the existing method, we recently introduced a new method that combines N-terminal bioconjugation methodology^28^ along with MAPLabTech^29^. This method indeed increases the diversity of proteins that can be used as scaffolds for the construction of SAPs. However, this method has three major limitations; (i) First, a vast number of proteins undergoes post-modification of the *N*-terminal group because of this reason, the above strategy would not work for those proteins (ii) secondly, in some cases, the N-terminal amine group may not be solvent-exposed and therefore not be available for the bioconjugation reaction (iii) finally, this method will not work if the protein contains proline in the second position^28^.

Site-specific labeling of a cysteine residue of a globular has been widely used for the synthesis of water-soluble non-self-assembling protein-polymer conjugates which include therapeutic proteins, antibody-drug conjugates. The ability to introduce cysteine moiety onto a protein at a predetermined position makes this methodology extremely powerful for a variety of applications. Although there are few reports on the synthesis of SAPs using chemical methods^30–35^, they have major limitations such as (i) they are not monodispersed (ii) bioconjugation reaction is mostly carried out in water/organic solvent mixture (iii) a viable and scalable method was not reported for purification. Therefore, it is important to develop a chemical methodology that addresses the above limitations. Towards that goal, herein, we report a chemical methodology for site-specific labeling of a cysteine residue of globular protein. The newly synthesized semi-synthetic was purified and characterized through various analytical techniques. Detailed studies show the designed protein self-assembled to form monodisperse protein nanoparticles. In addition, we also report the design, synthesis self-assembly, and disassembly studies of redox- and pH-responsive artificial proteins.

## Results

### Synthesis of Cysteine-Reactive amphiphilic activity-based probe

The probe contains three structural parts (i) maleimide targeting group (ii) precisely defined hydrophilic oligoethylene glycol and, (iii) hydrophobic tail **(Scheme 1)**. The choice of maleimide group is based on its selectivity towards cysteine residue in the presence of other potential functional groups such as primary amine, alcohol, etc. The choice of the chain length of oligoethylene glycol and hydrophobic tail length/branching was based on our extensive previous experience. Accordingly, we synthesized a maleimide probe **(MA-OEG-C18-1T)** through multi-step organic synthesis **(Scheme 1)**. In brief, hydrophilic alkyne (**1**) and hydrophobic azide (**2**) were allowed to react in the presence of 1M sodium ascorbate and 1M CuSO_4_ for 16 hrs at room temperature to get compound **3**. The amphiphilic tosylate (**3**) was treated with sodium azide to get compound **4**. The azide residue was subjected to reduction using triphenylphosphine to get compound 5 followed by treatment with N-(methoxycarbonyl) maleimide to get the final compound 6 **(MA-OEG-C18-1T)**.

### Bioconjugation Reaction and Self-assembly Studies

Bovine serum albumin (BSA) is chosen as a target protein following reasons (i) it is a globular proteins having a medium size and (ii) it contains one cysteine residue located on the outer surface of the protein. Triton X-100 was used to solubilize **MA-OEG-C18-1T** and the bioconjugation was carried out at room temperature in a 100% aqueous medium for 12 hours. The progress of the reaction was monitored using MALDI-TOF MS (Figure 1**a**). After 12 hours, we observed 70% conversion. After that reaction was over, the mixture was purified (Figure 1**b**) by ion-exchange chromatography (IEX) followed by size-exclusion chromatography (SEC). The self-assembling ability of the designed BSA conjugate was tested by analytical SEC (Figure 1**c**). As expected, the artificial BSA conjugate self-assemble to make protein nanoparticles of bigger size as evident from the elution profile seen in the SEC chromatogram. The relative molecule weight of protein nanoparticles was obtained from the standard calibration curve (Table 1). The obtained data suggest that the protein nanoparticles contain about 9-10 subunits of individual proteins. It is interesting to note that this conjugate self-assembled to form a slightly bigger complex than that of its corresponding N-terminus conjugate of the same spacer and tail group (Figure 1c, 1d and Table 1).

**Table 1.** Comparison of SEC data of N-terminus vs. thiol conjugate of BSA.

| S.No | Protein Conjugate | Elution Vol<br>SEC (mL) | RMW SEC<br>(kDa) | Oligomeric state<br>SEC (mer) | D<sub>h</sub> DLS<br>(nm) |
| --- | --- | --- | --- | --- | --- |
| 1 | BSA Nat | 17.4 | 66 | 1 | -- |
| 2 | BSA-OEG-C18-1T (N-Ter) | 13 | 700 | 10 | 13 |
| 2 | BSA-OEG-C18-1T (Thiol) | 13.8 | 630 | 9 | -- |

**Figure 1.**
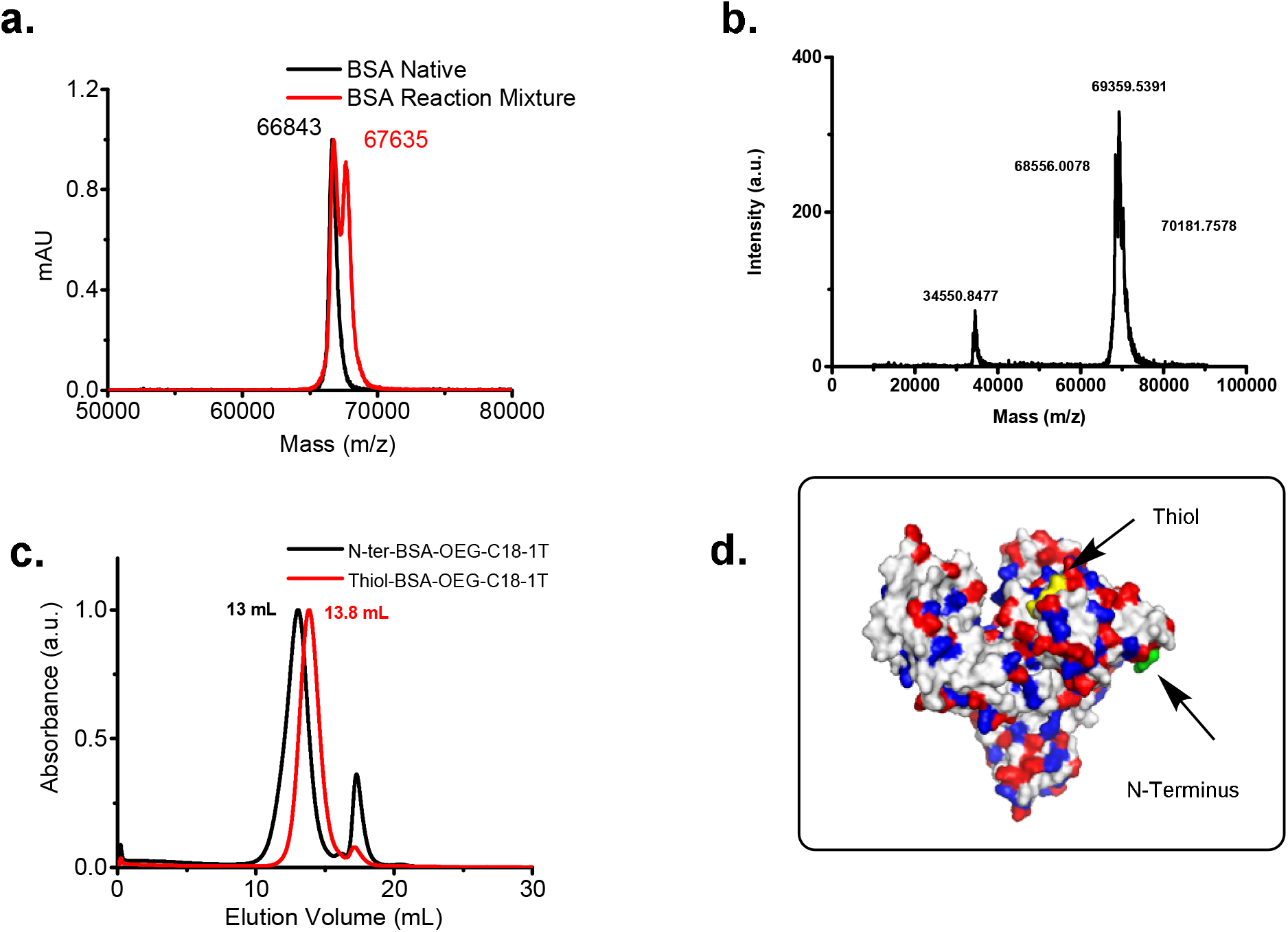
**a**. MALDI-TOF data of Native BSA and reaction mixture. **b**. MALDI-TOF data of purified BSA conjugate **c**. SEC chromatogram of BSA-OEG-C18-1T (Thiol) *vs* BSA-OEG-C18-1T (N-term). **d**. BSA (PDB accession: 3V03) structure showing thiol residue (yellow) with respect to N-terminus location (green). Negatively charged residues like Asp and Glu are colored red. Positively charged residues like Arg and Lys are colored blue.

### Molecular Design and Synthesis of redox-sensitive amphiphilic activity-based probe

Compound **7** was obtained by hydrolysis of compound **8** in sodium hydroxide. The obtained compound was allowed to react with 2, 2’-disulfanediylbis (ethan-1-ol) in presence of EDC and DMAP in DCM to afford compound **9**. This was followed by the activation using N N’-DSC, in the presence of triethyl amine to yield compound **10**. The activated ester **10** was then reacted with amine **(11)** in the presence of Et_3_N and DMF to obtain compound **12**. Then, the resultant diphosphonate ester **12** was heated with lithium bromide in DMF to get monophosphonate ester **13** which finally on fluorination using DAST in DCM afforded flurophosphonate **14 (FP-OEG-SS-C12-2T)**.

### Bioconjugation Reaction and Self-assembly/Disassembly Studies

The bioconjugation reaction, purification, and characterization of **Try-OEG-SS-C12-2T** were carried out similarly described in the previous section (Figure 2a and 2b). Analytical SEC results revealed that this conjugate can form protein nanoparticles as evident from elution volume (17.2 mL) in superpose 6 column and DLS (Figure 2c). The hydrodynamic diameter of the protein nanoparticle was determined using dynamic light scattering (DLS). As expected, the size of the protein nanoparticle is around 11 nm much bigger than the monomeric protein (Figure 2d). To get information on the molecular weight of the protein nanoparticles, we have carried out size-exclusion chromatography coupled with multi-angle light scattering (SEC-MALS) studies. The molecular weight of nanoparticles particle is around 425 kDa, from this data we have calculated the oligomeric state of the protein nanoparticles, which is around 17 sub-units. One of the most impressive features of the SEC-MALS data is that it gives information about the polydispersity of the protein nanoparticles. It is pretty evident from the data that these protein nanoparticles are extremely monodisperse (Figure 2e).

**Figure 2.**
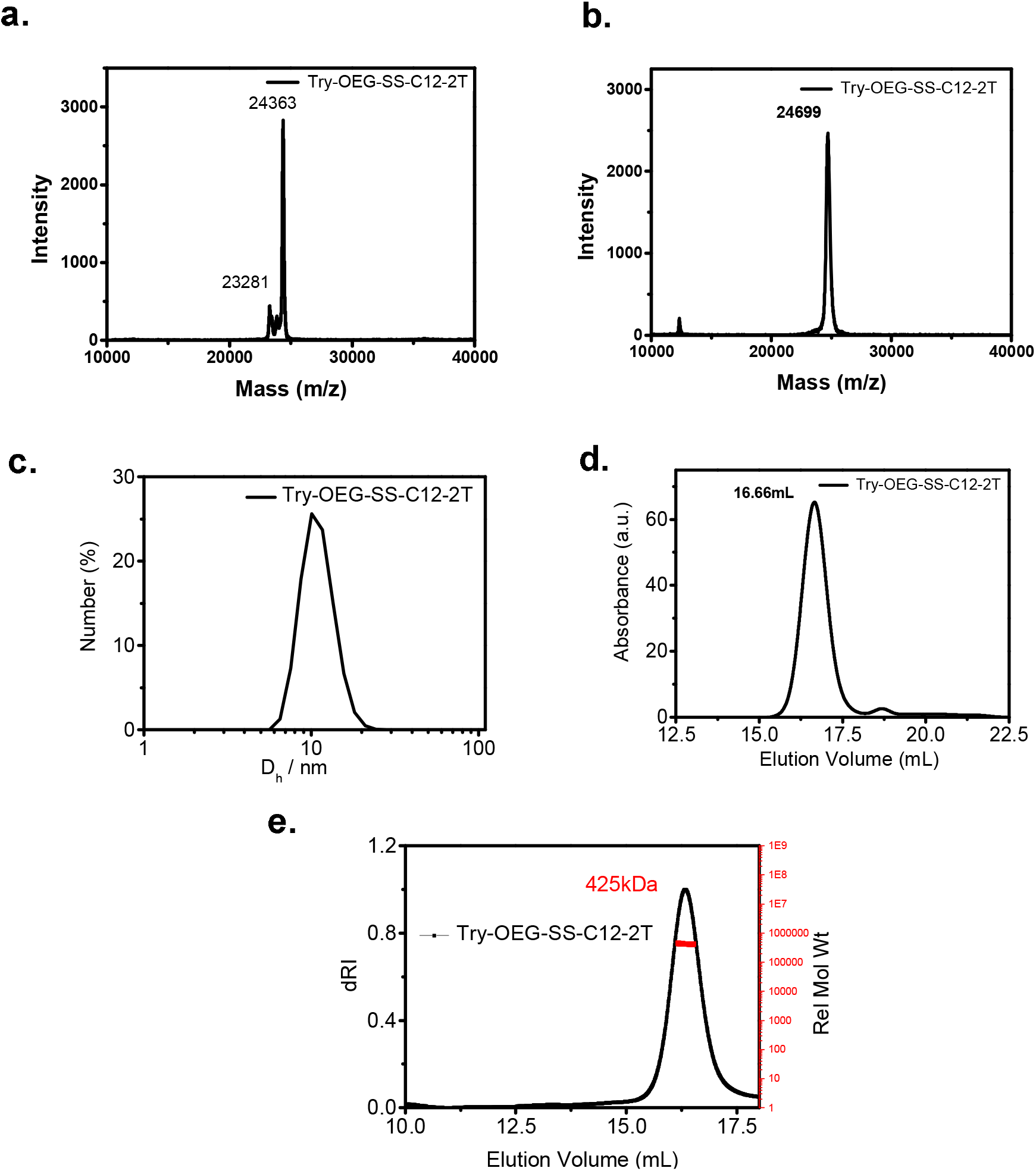
Characterization and self-assemby data for Try-OEG-SS-C12-2T. **a** MALDI-TOF. data of reaction mixture. **b**. MALDI-TOF data of purified conjugate. **c**. DLS data of purified conjugate. **d**. SEC chromatogram of purified conjugate. **e**. SEC-MALS data of purified conjugate.

After systematic self-assembly studies, we focused our attention on disassembly studies of protein nanoparticles **Try-OEG-SS-C12-2T**. We hypothesis that the selective cleave of “disulfide functionality” would convert self-assembling protein nanoparticles into non-self-assembling monomeric proteins. To test this hypothesis, we first carried out disassembly studies in the presence of different concentrations of DTT (Figure 3a). The addition of 10 and 20 equivalents of DTT resulted in the disassembly of protein nanoparticles to 80 to 90% as evident from the SEC results. However, the quantitative disassembly of protein nanoparticles was achieved using 30 equivalents of DTT (Figure 3a). Further, we wanted to check the kinetics of the disassembly reaction and therefore carried out the reactions at different time points in the presence of 30 equivalents of DTT. At 15 and 30 minutes, we observed about 80% and 90% disassembly. The quantitative conversion was achieved in 60 minutes (Figure 3b). Finally, we have confirmed the complete cleavage of **Try-OEG-SS-C12-2T** by using MALDI-TOF MS (Figure 3c).

**Figure 3.**
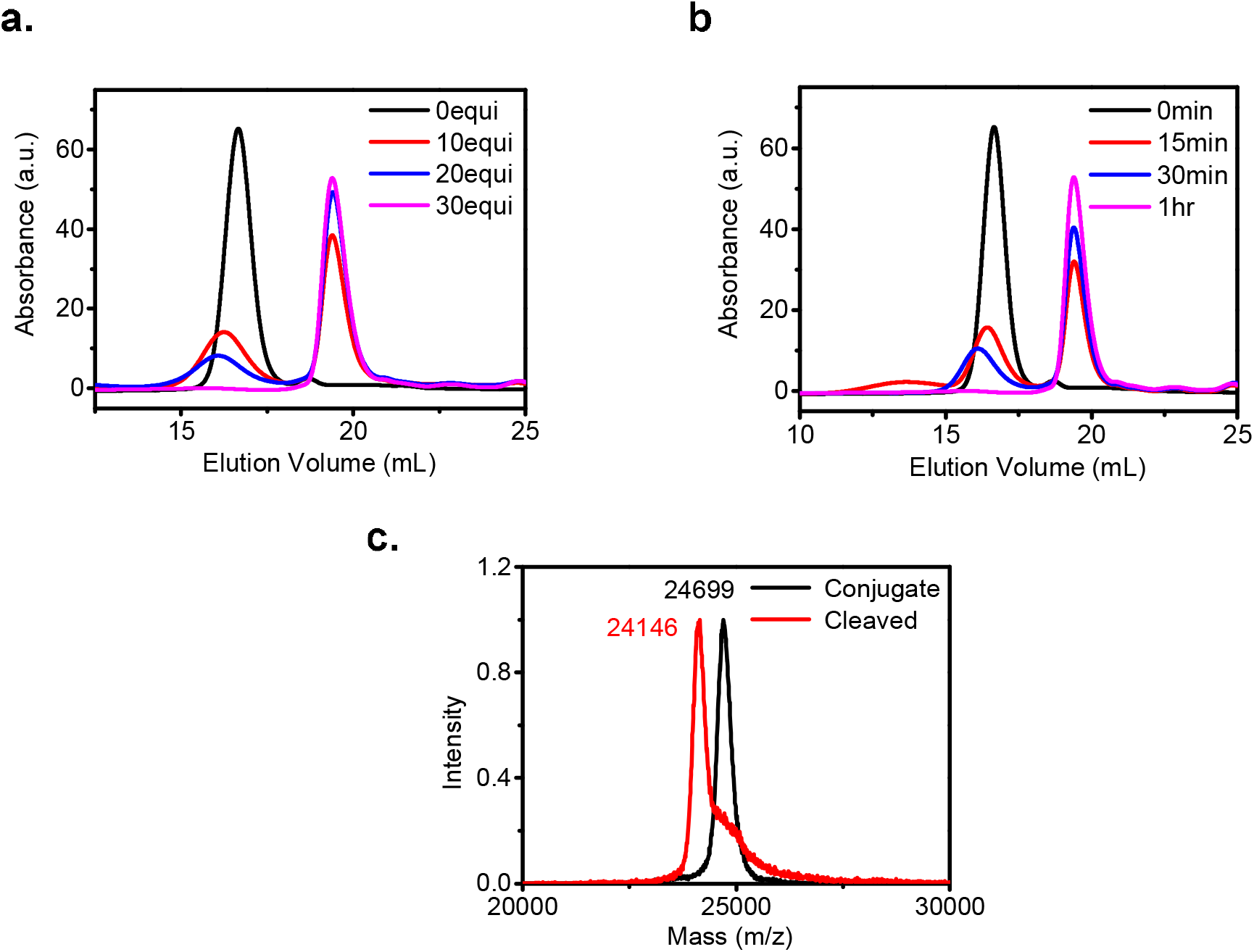
Dis-assembly data of Try-OEG-SS-C12-2T complex **a**. Dis-assembly studies at varying concentrations of DTT. **b**. Time-dependent disassembly studies **c**. MALDI-TOF spectra of purified (black) and cleaved conjugate (red).

To prove that the disassembly reaction is mediated by DTT, we have carried out the following control experiments. First, we tested the stability of the monomeric native protein in the presence of DTT, as expected, due to proteolysis the native protein was converted into small peptides as evident from the SEC results (Figure 4a). If the above result is true, then the active-site labeling of serine residue should block the proteolysis process and the protein should be intact. Indeed, we observed no cleave of monomeric protein when the active-site serine residue is blocked using an activity-based probe (Figure 4b). However, the modified protein is a monomeric and therefore not a one-to-one comparison. Therefore, we made another self-assembling artificial protein Tyr-OEG-C12-2T, this protein structure exactly mimics the structure of Tyr-OEG-SS-C12-2T, except the control protein, does not contain an engineered disulfide bond. More interestingly, both the proteins displayed the same elution profile in SEC (Figure 4c).

**Figure 4.**
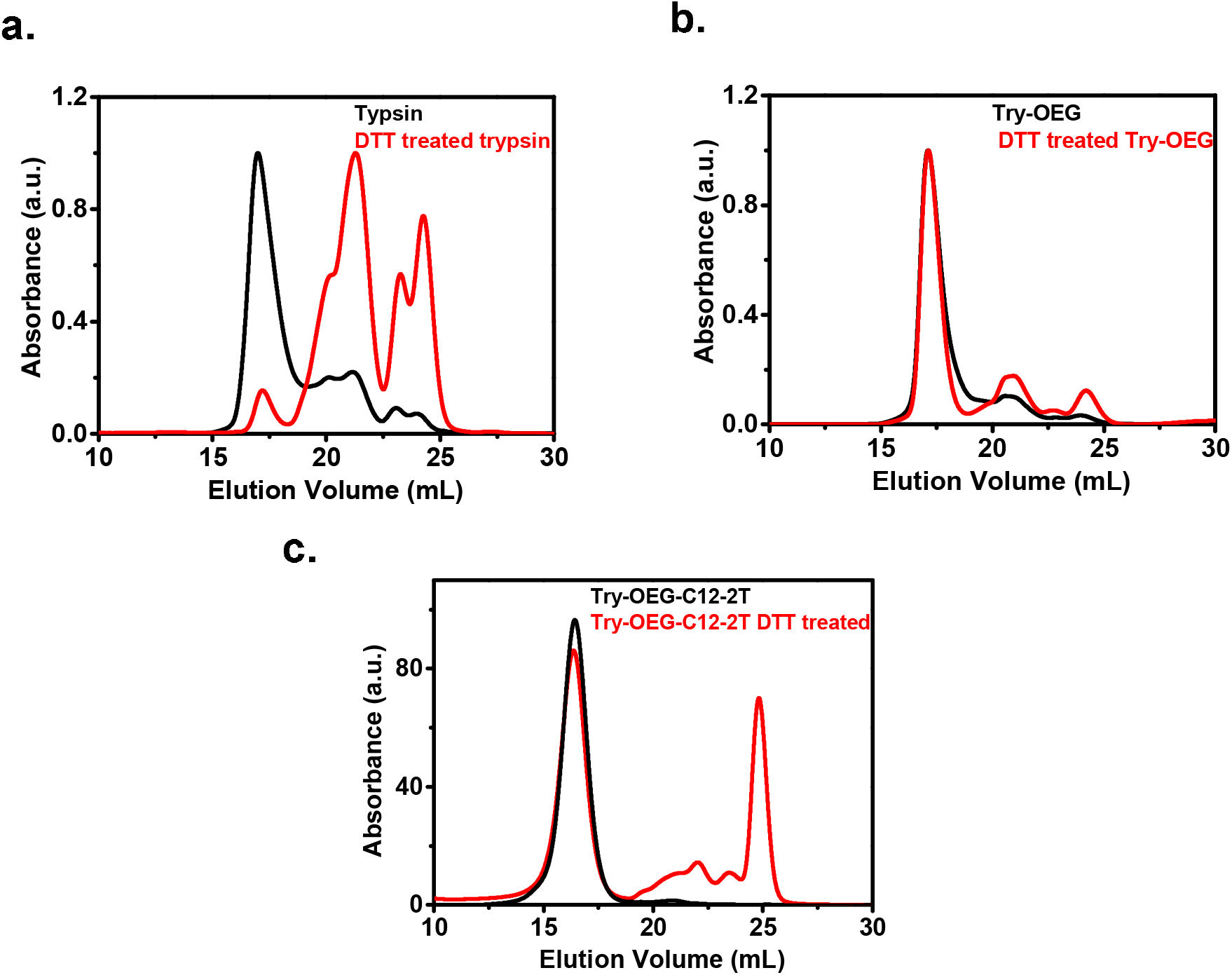
Control experiments **a**. SEC chromatogram of native trypsin (black) DTT treated trypsin (red). **b**. Try-OEG (black) and DTT treated try-OEG (red). **c**. Try-OEG-C12-2T (black) and DTT treated Try-OEG-C12-2T (red) respectively.

These results indicate that both SAPs can form protein nanoparticles of similar size. So, when we carried out the disassembly studies of Tyr-OEG-C12-2T, and Tyr-OEG-SS-C12-2T in the presence of DTT, Tyr-OEG-C12-2T did not disassemble whereas Tyr-OEG-SS-C12-2T disassembled into constitutive monomers as expected (Figure 3 and 4a). The above results strongly support our hypothesis that the disassembly reaction is driven by cleave of disulfide bond by the DTT reagent.

### Synthesis of a pH-Sensitive amphiphilic activity-based probe

Compound **17** was obtained from click reaction between azide **16** and alkyne **15** using CuSO_4_. Compound **17** was refluxed with hydrazine to yield compound **18**. The compound **19** upon refluxing with an aldehyde in ethanol resulted in diphosphonate ester compound **20**. Then, the resultant diphosphonate ester **20** was heated with sodium azide in DMF to get monophosphonate ester **21** which finally on fluorination using DAST in DCM afforded flurophosphonate **22 (FP-OEG-NN-C12-2T)**.

### Bioconjugation Reaction and Self-assembly/Disassembly Studies

For bioconjugation reaction, purification, and self-assembly studies, we followed the similar protocol described in the previous section (Figure 5a and 5b). Further, a disassembly reaction was carried out at pH 5 for several days. The SEC results suggest that there was a gradual decrease in the concentration of protein nanoparticles and a simultaneous increase in the concentration of monomeric protein (Figure 5c and 5d). However, we did not see observe quantitative conversion. After careful analysis, we figure out the presence of “imine functionality” next to the carbonyl group decreases the susceptibility of imine group hydrolysis at acidic pH. However, we need to do further experiments to support that hypothesis. However, those efforts are beyond the scope of the present work and therefore reported in the near future.

**Figure 5.**
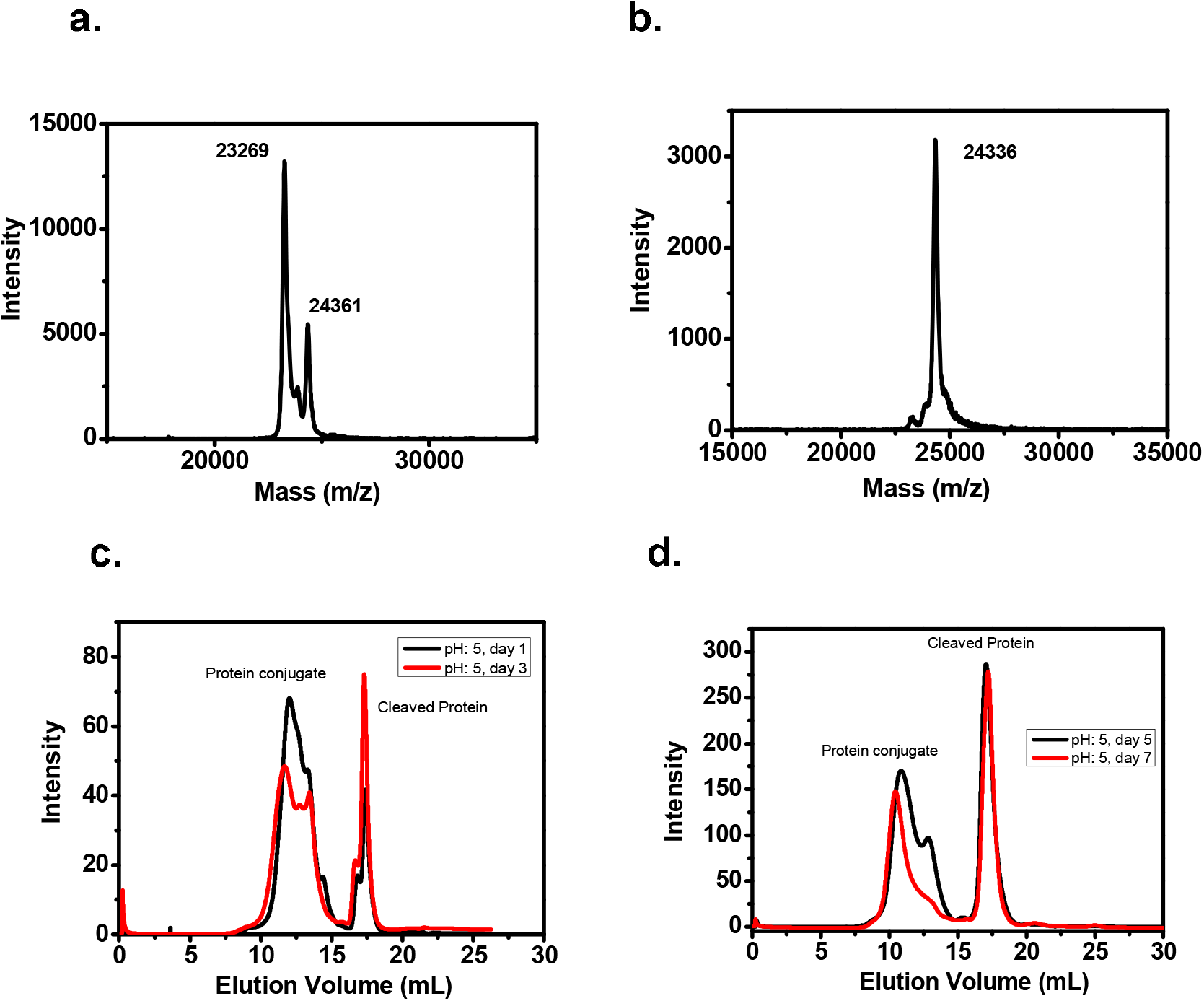
Characterization and pH-dependent disassembly studies. **a**. MALDI-TOF data of reaction mixture. **b**. MALDI-TOF data of purified conjugate. **e**. Dis-assembly studies on day 1 and day 2. **d**. Dis-assembly studies on day 5 and day 7.

## Discussion

Among the chemical technologies used for the design of well-defined monodisperse globular self-assembling artificial proteins, micelle-assisted protein labeling technology (MAPLabTech) has made enormous progress in the last 5 years^22–26^. The MAPLabTech 1.0 was effectively used for the synthesis of SAPs from the proteins belonging to the serine protease family. Further, MAPLab 2.0 was developed for labeling of *N*-terminal residue of any given protein which increased the scope of this technology substantially^29^. However, both methods have some limitations. MAPLabTech 1.0 can be only used for serine proteases. Similarly, MAPLabTech 2.0 does not apply proteins that are subjected to N-terminal post-translational modification, and most importantly the *N*-terminal amino acid should be solvent-exposed. Considering these limitations, we decided to add one more tool to the existing toolbox of MAPLabTech. Cysteine labeling is one of the few bioconjugation chemistries that have been widely used for various applications including synthesis for therapeutic purposes. Accordingly, we designed a cysteine-reactive amphiphilic probe and used MAPLabTech for labeling cysteine residue of BSA. As expected, modified BSA self-assembled to make protein nanoparticles of defined size with the precise oligomeric state. Although we have shown the applicability of this method with just one protein, in principle, it could be applied to any protein-containing natural or engineered cysteine.

We also designed and synthesized a redox-sensitive amphiphilic probe (RSAP), which was further used for the successful synthesis of SAPs containing an engineered disulfide bond. As expected, these proteins also self-assembled to make protein nanoparticles of defined size having the precise oligomeric state as evident from size-exclusion chromatography (SEC), dynamic light scattering (DLS), and size-exclusion chromatography coupled with multi-angle light scattering (SEC-MALS) results. The SEC-MALS data clear indicates these proteins nanoparticles are highly monodisperse and reminiscent of virus-like particles. The disassembly studies in presence of different concentrations of DTT revealed that 10 equivalents of DTT are enough to achieve complete disassembly. Time-dependent disassembly studies reveal that 1 hour is good enough to drive the disassembly reaction to completion. These results indicate that the artificial protein nanoparticles are quite porous and hence DTT reagent can access the disulfide functionality and thereby facilitate the cleavage of a disulfide bond.

As a further application, we designed and synthesized a pH-sensitive amphiphilic activity-based probe. This probe was successfully for bioconjugation reaction using MAPLabTech. The synthesized pH sensitive artificial protein self-assembled to make nanoparticles of defined as evident from SEC studies. However, disassembly studies at pH 5 revealed that the complexes are intact at acidic pH. The above results emphasize the importance of molecular design for achieving specific results in the field of biomedical science.

## Conclusion

We demonstrated in this study that the MAPLab technology can be effectively used for labeling surface-exposed cysteine residue. The synthesized artificial protein is self-assembled to form protein nanoparticles of defined size. Besides, the utility of MAPLab technology was demonstrated by the synthesis of redox- and pH-sensitive SAPs. Detailed self-assembly studies revealed that these proteins self-associate in a programmed way to make protein nanoparticles of defined size having a precise oligomeric state. Disassembly studies have shown that these nanoparticles can be disassembled upon applying a suitable redox trigger. Synthesis of pH-responsive protein amphiphile was also demonstrated nicely. As expected, they self-assemble to form nanoparticles of defined size. However, they fail to disassemble completely.

**Scheme 1.**
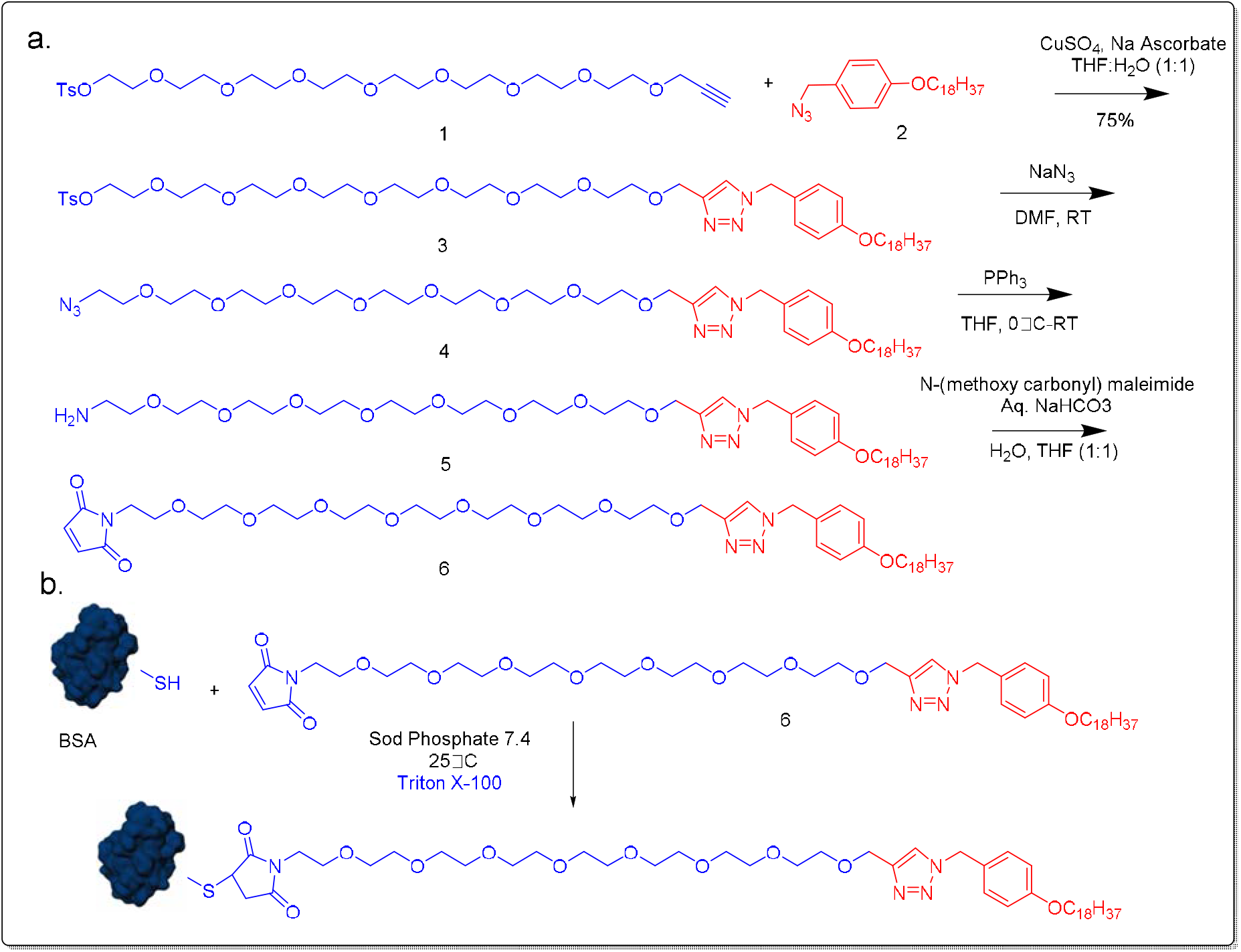
**a**. Scheme for synthesis of cysteine-reactive amphiphilic probe **b**. Bioconjugation reaction of BSA using MAPLabTech.

**Scheme 2.**
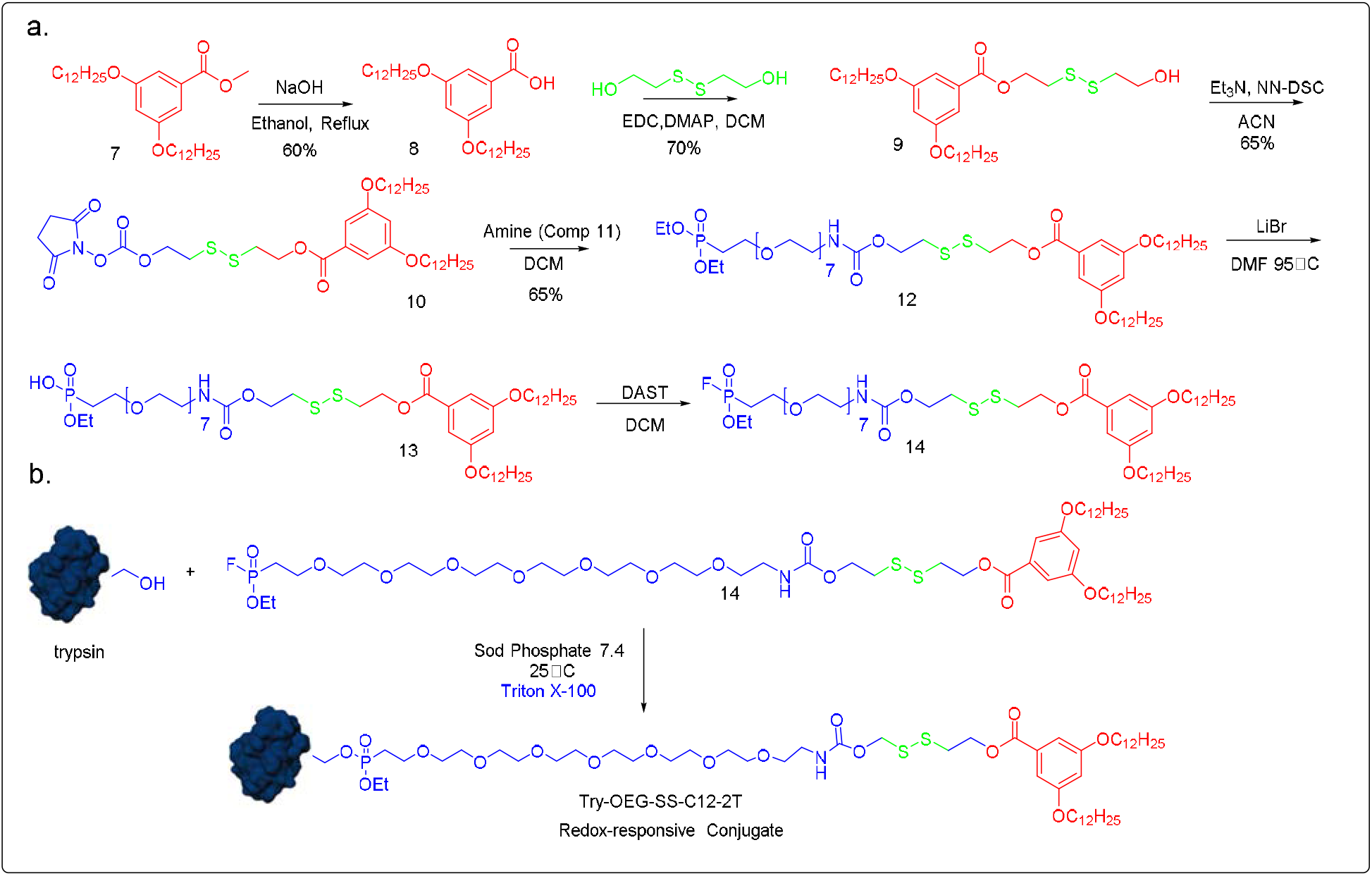
**a**. Synthesis of redox-sensitive amphiphilic activity-based probe. **b**. Bioconjugation reaction.

**Scheme 3.**
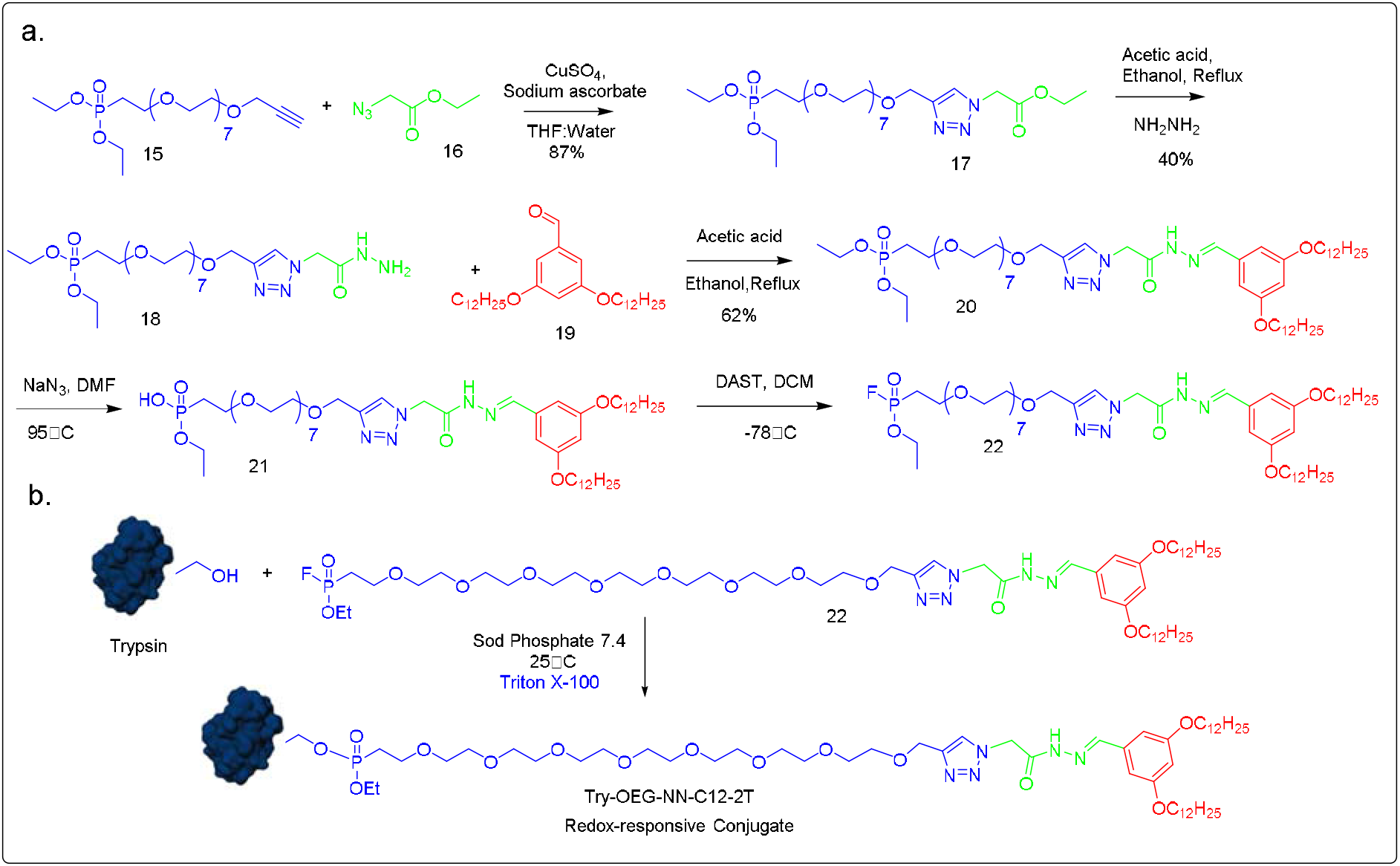
**a**. Scheme for synthesis of pH-sensitive activity-based amphiphilic probe **b**. Bioconjugation reaction.

**Scheme 4.**
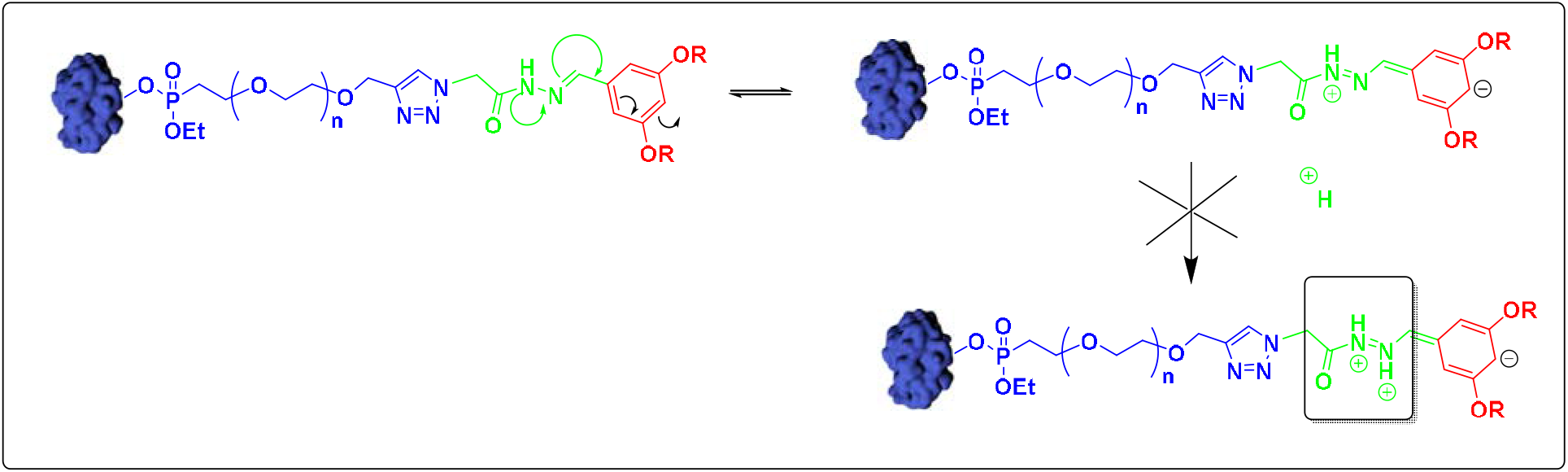
Proposed mechanism for pH-sensitive conjugate not getting cleaved.

## Associated Content

## Acknowledgments

This work was supported by the DST-SERB Early Career Award to B.S.S. DST-SERB (ECR/2015/000253) and the DBT grant (BT/PRI1450/BRB/I0/1370/2015).

## Conflict of Interest

We have filed a US patent for the technology disclosed in this paper

## TOC

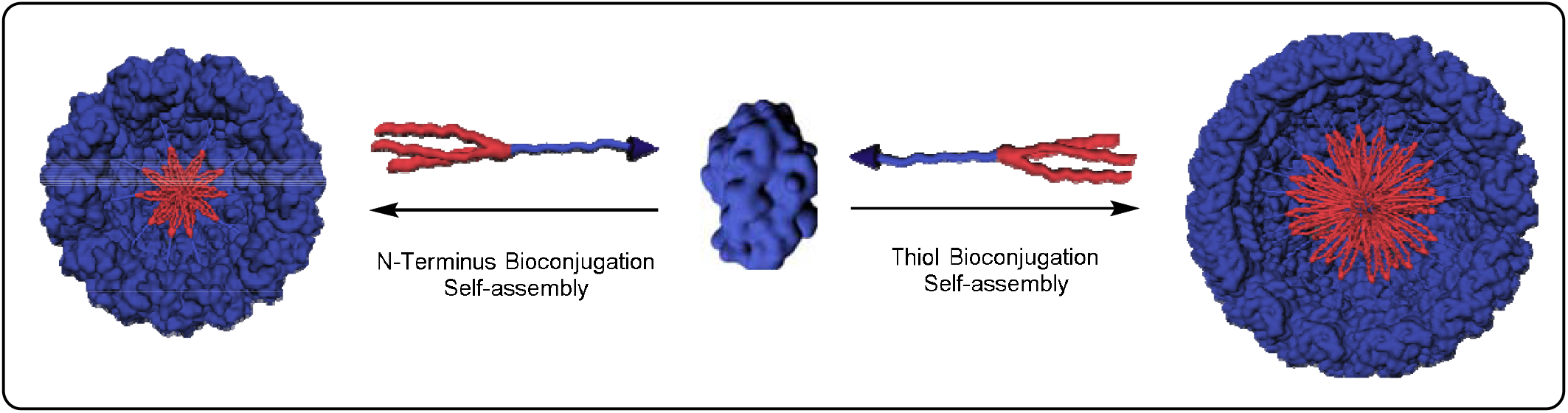

## References

(1) Huang, P. S.; Boyken, S. E.; Baker, D. The Coming of Age of de Novo Protein Design. Nature. 2016, 537, 320–327.

(2) Yeates, T. O. Geometric Principles for Designing Highly Symmetric Self-Assembling Protein Nanomaterials. Annu. Rev. Biophys. 2017, 46, 23–42.

(3) Reynhout, I. C.; Cornelissen, J. J. L. M.; Nolte, R. J. M. Synthesis of Polymer-Biohybrids: From Small to Giant Surfactants. Acc. Chem. Res. 2009, 42 (6), 681–692.

(4) Luo, Q.; Hou, C.; Bai, Y.; Wang, R.; Liu, J. Protein Assembly: Versatile Approaches to Construct Highly Ordered Nanostructures. Chem. Rev. 2016, 116, 13571–13632.

(5) Luo, Q.; Dong, Z.; Hou, C.; Liu, J. A One-Pot, Three-Component Reaction for the Synthesis of Novel 7-Arylbenzo[c]Acridine-5,6-Diones. Chem. Commun. 2014, 50, 9997– 10007.

(6) Purwar, M.; Pokorski, J. K.; Singh, P.; Bhattacharyya, S.; Arendt, H.; DeStefano, J.; La Branche, C. C.; Montefiori, D. C.; Finn, M. G.; Varadarajan, R. Design, Display and Immunogenicity of HIV1 Gp120 Fragment Immunogens on Virus-like Particles. Vaccine 2018, 36, 6345–6353.

(7) Smith, M. L.; Lindbo, J. A.; Dillard-Telm, S.; Brosio, P. M.; Lasnik, A. B.; McCormick, A. A.; Nguyen, L. V.; Palmer, K. E. Modified Tobacco Mosaic Virus Particles as Scaffolds for Display of Protein Antigens for Vaccine Applications. Virology 2006, 348, 475–488.

(8) Ma, Y.; Nolte, R. J. M.; Cornelissen, J. J. L. M. Virus-Based Nanocarriers for Drug Delivery. Adv. Drug Deliv. Rev. 2012, 64, 811–825.

(9) Weissleder, R.; Nahrendorf, M.; Pittet, M. J. Imaging Macrophages with Nanoparticles. Nat. Mater. 2014, 13, 125–138.

(10) Brunel, F. M.; Lewis, J. D.; Destito, G.; Steinmetz, N. F.; Manchester, M.; Stuhlmann, H.; Dawson, P. E. Hydrazone Ligation Strategy to Assemble Multifunctional Viral Nanoparticles for Cell Imaging and Tumor Targeting. Nano Lett. 2010, 10, 1093–1097.

(11) Nagaoka, M.; Jiang, H. L.; Hoshiba, T.; Akaike, T.; Cho, C. S. Application of Recombinant Fusion Proteins for Tissue Engineering. In Annals of Biomedical Engineering 2010, 38, 683–893

(12) McGann, C. L.; Levenson, E. A.; Kiick, K. L. Resilin-Based Hybrid Hydrogels for Cardiovascular Tissue Engineering. Macromol. Chem. Phys. 2013, 214, 203–213

(13) Hsia, Y.; Bale, J. B.; Gonen, S.; Shi, D.; Sheffler, W.; Fong, K. K.; Nattermann, U.; Xu, C.; Huang, P. S.; Ravichandran, R.; Yi, S.; Davis, T. N.; Gonen, T.; King, N. P.; Baker, D. Design of a Hyperstable 60-Subunit Protein Icosahedron. Nature 2016, 535, 136–139.

(14) Bale, J. B.; Gonen, S.; Liu, Y.; Sheffler, W.; Ellis, D.; Thomas, C.; Cascio, D.; Yeates, T. O.; Gonen, T.; King, N. P.; Baker, D. Accurate Design of Megadalton-Scale Two-Component Icosahedral Protein Complexes. Science 2016, 353, 389–394.

(15) Brouwer, P. J. M.; Antanasijevic, A.; Berndsen, Z.; Yasmeen, A.; Fiala, B.; Bijl, T. P. L.; Bontjer, I.; Bale, J. B.; Sheffler, W.; Allen, J. D.; Schorcht, A.; Burger, J. A.; Camacho, M.; Ellis, D.; Cottrell, C. A.; Behrens, A. J.; Catalano, M.; del Moral-Sánchez, I.; Ketas, T. J.; LaBranche, C.; van Gils, M. J.; Sliepen, K.; Stewart, L. J.; Crispin, M.; Montefiori, D. C.; Baker, D.; Moore, J. P.; Klasse, P. J.; Ward, A. B.; King, N. P.; Sanders, R. W. Enhancing and Shaping the Immunogenicity of Native-like HIV-1 Envelope Trimers with a Two-Component Protein Nanoparticle. Nat. Commun. 2019, 10,

(16) King, N. P.; Sheffler, W.; Sawaya, M. R.; Vollmar, B. S.; Sumida, J. P.; André, I.; Gonen, T.; Yeates, T. O.; Baker, D. Computational Design of Self-Assembling Protein Nanomaterials with Atomic Level Accuracy. Science 2012, 336, 1171–1174.

(17) Padilla, J. E.; Colovos, C.; Yeates, T. O. Nanohedra: Using Symmetry to Design Self Assembling Protein Cages, Layers, Crystals, and Filaments. Proc. Natl. Acad. Sci. U. S. A. 2001, 98, 2217–2221.

(18) Lai, Y. T.; Reading, E.; Hura, G. L.; Tsai, K. L.; Laganowsky, A.; Asturias, F. J.; Tainer, J. A.; Robinson, C. V.; Yeates, T. O. Structure of a Designed Protein Cage That Self-Assembles into a Highly Porous Cube. Nat. Chem. 2014, 6, 1065–1071.

(19) Kobayashi, N.; Yanase, K.; Sato, T.; Unzai, S.; Hecht, M. H.; Arai, R. Self-Assembling Nano-Architectures Created from a Protein Nano-Building Block Using an Intermolecularly Folded Dimeric de Novo Protein. J. Am. Chem. Soc. 2015, 137, 11285– 11293.

(20) Tetter, S.; Terasaka, N.; Steinauer, A.; Bingham, R. J.; Clark, S.; Scott, A. J. P.; Patel, N.; Leibundgut, M.; Wroblewski, E.; Ban, N.; Stockley, P. G.; Twarock, R.; Hilvert, D. Evolution of a Virus-like Architecture and Packaging Mechanism in a Repurposed Bacterial Protein. Science 2021, 372, 6547.

(21) Mozhdehi, D.; Luginbuhl, K. M.; Dzuricky, M.; Costa, S. A.; Xiong, S.; Huang, F. C.; Lewis, M. M.; Zelenetz, S. R.; Colby, C. D.; Chilkoti, A. Genetically Encoded Cholesterol-Modified Polypeptides. J. Am. Chem. Soc. 2019, 141, 945–951.

(22) Sandanaraj, B. S.; Reddy, M. M.; Bhandari, P. J.; Kumar, S.; Aswal, V. K. Rational Design of Supramolecular Dynamic Protein Assemblies by Using a Micelle-Assisted Activity-Based Protein-Labeling Technology. Chem. - A Eur. J. 2018, 24, 16085–16096.

(23) Sandanaraj, B.; Bhandari, P. J.; Reddy, M. M.; Lohote, A. B.; Sahoo, B. Design, Synthesis and Self_jassembly Studies of Suite of Monodisperse Facially Amphiphilic Protein_jDendron Conjugates. ChemBioChem 2019, 21, 408–416

(24) Sandanaraj, B. S.; Reddy, M. M.; Rao, K. J.; Bhandari, P. J. Rational Design of Semi-Synthetic Protein Complexes with the Defined Oligomeric State. ChemistrySelect 2019, 4 (20), 6397–6402.

(25) Bhandari, P. J.; Sandanaraj, B. S. Programmed and Sequential Disassembly of Multi-Responsive Supramolecular Protein Nanoassemblies: A Detailed Mechanistic Investigation. ChemBioChem 2021, 22, 876–887.

(26) Bhandari, P. J.; Sandanaraj, B. S. Rational Design of Programmable Monodisperse Semi-Synthetic Protein Nanomaterials Containing Engineered Disulfide Functionality. 2021. https://doi.org/10.1002/cbic.202100288

(27) Junutula, J. R.; Raab, H.; Clark, S.; Bhakta, S.; Leipold, D. D.; Weir, S.; Chen, Y.; Simpson, M.; Tsai, S. P.; Dennis, M. S.; Lu, Y.; Meng, Y. G.; Ng, C.; Yang, J.; Lee, C. C.; Duenas, E.; Gorrell, J.; Katta, V.; Kim, A.; McDorman, K.; Flagella, K.; Venook, R.; Ross, S.; Spencer, S. D.; Lee Wong, W.; Lowman, H. B.; Vandlen, R.; Sliwkowski, M. X.; Scheller, R. H.; Polakis, P.; Mallet, W. Site-Specific Conjugation of a Cytotoxic Drug to an Antibody Improves the Therapeutic Index. Nat. Biotechnol. 2008, 26, 925–932.

(28) Macdonald, J. I.; Munch, H. K.; Moore, T.; Francis, M. B. One-Step Site-Specific Modification of Native Proteins with 2-Pyridinecarboxyaldehydes. Nat. Chem. Biol. 2015, 11, 326–331.

(29) (a) Reddy, M. M.; Bathla, P.; Sandanaraj, B. S. A Universal Chemical Method for Rational Design of Protein-Based Nanoreactors. bioRxiv 2021, 2021.03.01.433350. (b) Reddy, M. M.; Bathla, P.; Sandanaraj, B. S. A Universal Chemical Method for Rational Design of Protein-Based Nanoreactors. ChemBioChem 2021 Accepted Article

(30) Velonia, K.; Rowan, A. E.; Nolte, R. J. M. Lipase Polystyrene Giant Amphiphiles. J. Am. Chem. Soc. 2002, 124, 4224–4225.

(31) Dirks, A. J.; Van Berkel, S. S.; Hatzakis, N. S.; Opsteen, J. A.; Van Delft, F. L.; Cornelissen, J. J. L. M.; Rowan, A. E.; Van Hest, J. C. M.; Rutjes, F. P. J. T.; Nolte, R. J. M. Preparation of Biohybrid Amphiphiles via the Copper Catalysed Huisgen [3 + 2] Dipolar Cycloaddition Reaction. Chem. Commun. 2005, 33, 4172–4174.

(32) Dirks, A. J.; Nolte, R. J. M.; Cornelissen, J. J. L. M. Protein-Polymer Hybrid Amphiphiles. Adv. Mater. 2008, 20, 3953–3957.

(33) Thomas, C. S.; Glassman, M. J.; Olsen, B. D.; Al, T. E. T. Solid-State Nanostructured Materials from Self-Assembly of a Globular Protein À Polymer Diblock Copolymer. 2011, 7, 5697–5707

(34) Thomas, C. S.; Xu, L.; Olsen, B. D. Kinetically Controlled Nanostructure Formation in Self-Assembled Globular Protein-Polymer Diblock Copolymers. Biomacromolecules 2012, 13, 2781–2792.

(35) Heredia, K. L.; Bontempo, D.; Ly, T.; Byers, J. T.; Halstenberg, S.; Maynard, H. D. In Situ Preparation of Protein - “Smart” Polymer Conjugates with Retention of Bioactivity. J. Am. Chem. Soc. 2005, 127, 16955–16960.

