## Supporting Information for "Rational Design of Self-assembling Artificial Proteins Utilizing a Micelle-Assisted Protein Labeling Technology (*MAPLabTech*): Testing the Scope"

**Experimental section and methods**

**Matrix preparation and molecular weight determination**

Apart from the molecular weight determination of native proteins, all stages of protein modification and purification were also followed by MALDI-TOF MS. The samples were analyzed in Linear High Mass mode in AB Sciex 4800 plus MALDI-TOF/TOF analyzer with 4000 Series Explorer as software. Mass was scanned between 10,000 Da and 90,000 Da with a focus mass, selected depending on the protein analyzed.

For trypsin the procedure remains same as our reported protocol. For BSA, we followed a different MALDI-TOF Ms Procedure in this work. To brief, 15mg of sinapinic acid was weighed in microcentrifuge tube then add 1.0 ml of matrix solution (70:30 water/acetonitrile with 0.1 % TFA final concentration) was added and vortexed to get the matrix mixture. The sample and matrix mixture were mixed at a 1:10 ratio. When the crystallization observed in centrifuge tubes, 1-2µL of the samples were spotted on the plate and air-dried for 15 minutes. Then the plate was loaded and fired to get accurate molecular weights.

**Protein modification, monitoring, and purification**

Trypsin modification, monitoring, and purification were done using a similar protocol given in our previous work. However, some changes were made in the equivalents we used and triton X-100 concentrations we used in the case of BSA. The AABP was used varied between 25 - 100 eq, which was just 2 eq in our previous work. Triton X-100, which was used concentrations 100 times (20 mM) more than CMC or 2% of the total volume of the reaction mixture to solubilize the AABPs in the previous work, gave noisy MALDI-TOF Spectra. So we decreased the Triton X-100 concentration to 10 times more than CMC. Heating the reaction mixture at 37⁰C triggered precipitation of protein conjugates in the reaction mixtures. So even though we see increased conversions with heating, we did the conjugation at RT.

All the conjugates were purified by two-step purification, *i.e.,* IEX and SEC, performed using FPLC. IEX was performed to remove triton X-100 using either SP sepharose or Q sepharose resins (GE) depending on isoelectric point (pI) and surface charges of proteins. For example, to purify the reaction mixture of BSA (pI: 4.7) we used Q sepharose, an anion-exchange resin at pH 7.4. The obtained IEX fractions were subjected to SEC immediately to remove the native proteins from the protein conjugates in 50 mM sodium phosphate pH 7.4, 1 M NaCl using either Sephacryl S-100 HR 16/60 or Sephacryl S-200 HR 16/60 or Sephacryl S-300 HR 16/60. After SEC the samples were stored at -80 ⁰C

**Synthesis and purification of BSA-Thiol conjugate**

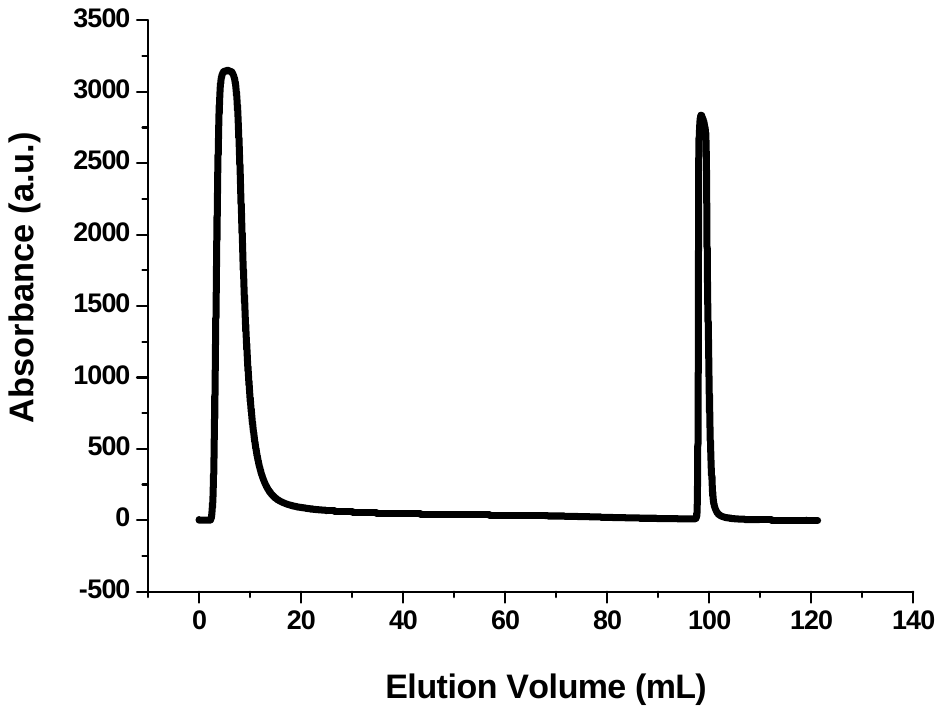

**Figure 4E.** IEX chromatogram of BSA-Thiol conjugation.

**Size exclusion-based molecular weight determination**

To determine the molecular weights of trypsin protein complexes, we performed SEC in Superdex 200 (GE healthcare). For BSA protein complexes, we performed SEC in Superose 6 (GE healthcare).

A series of size exclusion runs were performed for standard proteins (GE Healthcare), in 50 mM sodium phosphate pH 7.4, 200 mM NaCl with 0.25 mL/min as the flow rates in respective columns. The calibration curve was plotted for K_av_ of standards against relative molecular weights (RMW) of standard proteins. The calibration plots were given in our previous reports.

**Other characterizations**

DLS, SEC-MALS were done in the same fashion using the same instruments given in our previous work.

**Synthesis and purification of AABPs and their intermediates**

All compounds were made, characterized using similar conditions and instruments as mentioned in our previous reports.

**Synthesis of maleimide** **AABP and its intermediates**

**Synthesis of Compound 4**

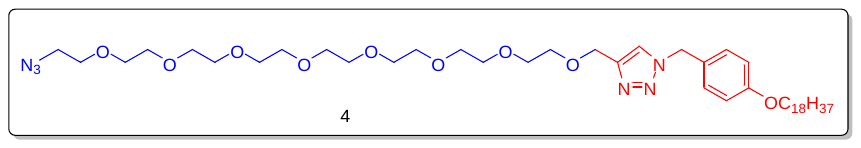

Mol. formula:	C_44_H_78_N_6_O_9_

Mol. weight: 835.14 g/mol

Physical appearance: yellow waxy solid

Yield: 70 %).

Tosylate (**compound 3**) (2 g, 2.0 mmol) was dissolved in DMF (10 mL) at RT. Then NaN_3_ (0.674 g, 10.3 mmol) was added to the reaction mixture and allowed to react at RT for 12 hrs. After this, the reaction mixture was concentrated and purified by NPC without any workup (1.2 g, 70 %). **^1^H NMR** (400 MHz, CDCl_3_): δ_H_ 7.40 (s, 1H), 7.20 (d, *J* = 8 Hz, 2H), 6.86 (d, *J* = 8 Hz, 2H), 5.41 (s, 2H), 4.62 (s, 2H), 3.91 (t, *J* = 6.8 Hz, 2H), 3.63 (m, 32H), 3.35 (t, *J* = 4.8 Hz, 2H), 1.74 (t, *J* = 6.8 Hz, 2H), 1.42 (m, 2H), 1.2 (m, 32H), 0.85 (t, *J* = 6.4, 3H). **MALDI-TOF MS** (MW+ Na): 874.58 g/mol.

**Synthesis of Compound 5**

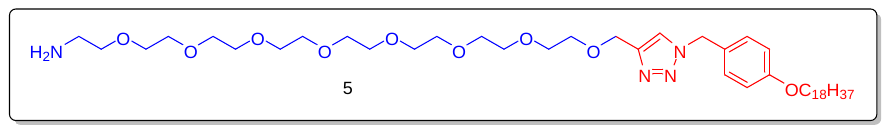

Mol. formula:	C_44_H_80_N_4_O_9_

Mol. weight: 808.14 g/mol

Physical appearance: yellow waxy solid

Yield: 51 %

Azide (**compound 4**) (400 mg, 0.479 mmol) was dissolved in THF (7 mL) at 0 ⁰C and PPh_3_ (250 mg, 0.954 mmol) dissolved in THF (3 mL) and was added at 0 ⁰C. Then the reaction mixture was allowed to react at RT. Water was then added to the reaction mixture after 12 hrs and allowed to stir for another 1 hr. Then the reaction mixture was extracted with DCM and purified using NPC (200 mg, 51 %). **^1^H NMR** (400 MHz, CDCl_3_): δ_H_ 7.40 (s, 1H), 7.20 (d, *J* = 8.8 Hz, 2H), 6.86 (d, *J* = 8.8 Hz, 2H), 5.43 (s, 2H), 4.64 (s, 2H), 3.93 (t, *J* = 6.8 Hz, 2H), 3.68 (m, 32H), 3.38 (t, *J* = 4.8 Hz, 2H), 1.76 (m, 2H), 1.42 (m, 2H), 1.33 (m, 30H), 0.87 (t, *J* = 7.2 Hz, 3H). **MALDI-TOF MS** (MW+ Na): 831.53 g/mol.

**Synthesis of Compound 6**

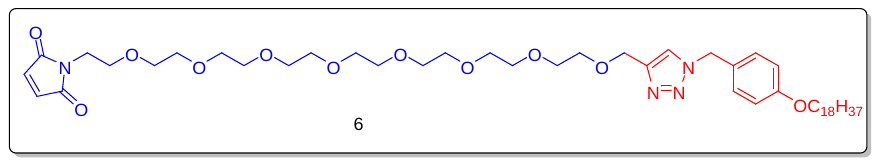

Mol. formula:	C_48_H_80_N_4_O_11_

Mol. weight: 889.58 g/mol

Physical appearance: yellow waxy solid

Yield: 50 %

Amine (**compound 5)** (100 mg, 0.123 mmol) was dissolved in a mixture of saturated aqueous solution of NaHCO_3_ (5 ml) and THF (5 ml) cooled on an ice bath. Then N-(methoxy carbonyl) maleimide (23 mg, 0.148 mmol) was added in portions over 5 mins under vigorous stirring. The mixture was stirred for 1 hr at 0 ⁰C, followed by 1 hr at room temperature. After extraction with DCM, the organic phase was dried over anhydrous Na_2_SO_4_, ﬁltered, and concentrated. Puriﬁcation by silica gel column chromatography (MeOH/DCM) yielded the product as colorless oil after evaporation with DCM (55 mg, 50 %). **^1^H NMR** (400 MHz, CDCl_3_): δ_H_ 7.43 (s, 1H), 7.20 (d, *J* = 8.8 Hz, 2H), 6.86 (d, *J* = 8.8 Hz, 2H), 6.68 (s, 2H), 5.46 (s, 2H), 4.62 (s, 2H), 3.91 (t, *J* = 6.1 Hz, 2H), 3.70 (t, *J* = 5.6 Hz, 2H), 3.63 (m, 32H), 1.74 (m, 2H), 1.41 (m, 2H), 1.23 (m, 32H), 0.85 (t, *J* = 7.2 Hz, 3H). **MALDI-TOF MS** (MW+ Na): 911.58 g/mol.

**Synthesis of redox-responsive** **AABP and its intermediates**

**Synthesis of compound 8**

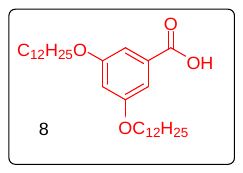

Mol. formula: C_31_H_54_O_4_

Mol. Weight: 490.76 g/mol

Physical appearance: White solid

Yield: 60 %

The compound (**7**) (3 g, 5 mmol), sodium hydroxide (NaOH) (0.5 g, 10 mmol) were dissolved in EtOH. The mixture was then refluxed for 4 hrs. Upon completion, the reaction was quenched with dropwise addition of water and acidified with conc. HCl. The obtained precipitate was filtered and washed with EtOH for two times. The combined organic layer dried over Na_2_SO_4_ and concentrated under reduced pressure to get the crude product, which was utilized for the next reaction without further purification. The product was obtained as white solid (1.7 g, 3 mmol, 60 %), Rf = 0.10 in 10 % EtOAc /PE. **^1^H NMR** (400 MHz, CDCl_3_): δ_H_ 7.21 (d, *J* = 2.4 Hz, 2H), 6.68 (t, *J* = 2.4 Hz, 1H), 3.98 (t, *J* = 6.4 Hz, 2H), 1.85-1.73 (m, 4H), 1.52-1.17 (m, 38H), 0.88 (t, *J* = 6.8 Hz ,6H).

**Synthesis of compound 9**

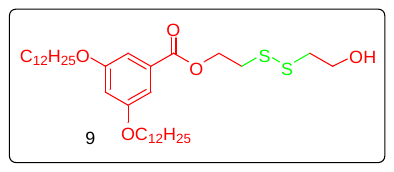

Mol. formula: C_35_H_62_O_5_S_2_

Mol. Weight: 626.99 g/mol

Physical appearance: White solid

Yield: 70 %

In an oven dried RBF **compound 8** (2.5 g, 5 mmol), 2,2'-disulfanediylbis(ethan-1-ol) (1.9 g, 10 mmol) and DMAP (0.3 g, 2 mmol) were taken and dissolved in DCM under stirring. Then, solution of EDC in DCM (2 g, 10 mmol) was added slowly to above mixture and allowed to stir for 12 hrs. Upon completion, reaction was quenched with water and resulting content was extracted in DCM thrice. Combined organic layers was dried over Na_2_SO_4_ and concentrated under reduced pressure to get crude product which was purified using silica gel column chromatography. The product (**9**) obtained was white solid (2.2 g, 3 mmol, 70 %), Rf = 0.20 in 10 % EtOAc / PE. **^1^H NMR** (400 MHz, CDCl_3_): δ_H_ 7.16 (d, *J* = 2.4 Hz, 2H), 6.64 (t, *J* = 2.4 Hz, 1H), 4.57 (t, *J* = 6.4 Hz, 2H), 3.96 (t, *J* = 6.4 Hz, 2H), 3.91-9.87 (m, 2H),3.05 (t, *J* = 6.8 Hz, 2H), 2.90 (t, *J* = 6.8 Hz, 2H), 1.84-1.73 (m, 4H), 1.52-1.26 (m, 38H), 0.88 (t, *J* = 6.8 Hz, 6H). **^13^C NMR** (100 MHz, CDCl_3_): δ_C_ 166.48, 160.30, 131.58, 107.88, 106.76, 77.16, 68.47, 63.12,60.33, 41.81, 37.12, 32.05, 29.79, 29.77, 29.73, 29.71, 29.49, 29.48, 29.30, 26.14, 22.82, 14.25**. MALDI-TOF MS** (M+Na): 665.47 g/mol.

**Synthesis of compound 10**

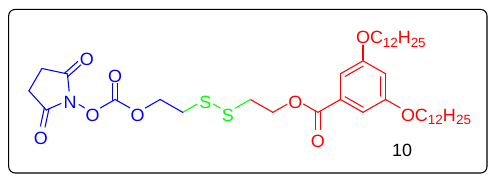

Mol. formula: C_40_H_65_NO_9_S_2_

Mol. Weight: 768.07 g/mol

Physical appearance: White solid

Yield: 65 %

The **compound** **10** was prepared from **compound** **9** (0.5 g, 0.8 mmol) and N,N′-DSC (1.6 g, 6 mmol) and Et_3_N (0.6 g, 5 mmol). The product obtained was pale yellow solid (0.4 g, 0.5 mmol, 65 %), Rf = 0.18 in 25 % EtOAc / PE. **^1^H NMR** (400 MHz, CDCl_3_): δ_H_ 7.14 (d, *J* = 2.4 Hz, 2H), 6.63 (t, *J* = 2.4 Hz, 1H), 4.59-4.54 (m, 4H), 3.96 (t, *J* = 6.4 Hz, 2H), 3.08-3.01 (m, 4H), 2.82(s, 4H), 1.80-1.73 (m, 4H), 1.47-1.26 (m, 38H), 0.87 (t, *J* = 6.8 Hz ,6H). **^13^C NMR** (100 MHz, CDCl_3_): δ_C_ 168.60, 166.33, 160.28, 151.54, 131.60, 107.85, 106.75, 68.84, 68.45, 62.96, 37.32, 36.39, 32.03, 29.78, 29.75, 29.72, 29.69, 29.49, 29.47, 29.29, 26.13, 25.56, 22.80, 14.24.

**Synthesis of compound 12**

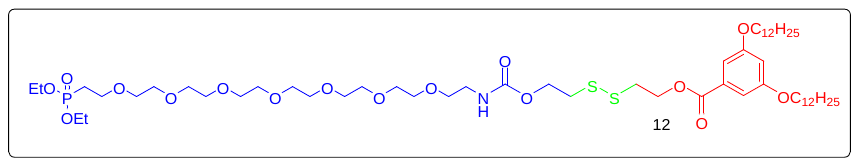

Mol. formula: C_56_H_104_NO_10_PS_2_

Mol. Weight: 1142.53 g/mol

Physical appearance: White solid

Yield: 65 %

The **compound** **12** was prepared from **compound** **10** (0.10 g, 0.13 mmol) and **amine (compound** **11)** (0.06 g, 0.11 mmol). The product obtained was pale yellow liquid (0.09 g, 0.07 mmol, 65 %), Rf = 0.40 in 5 % MeOH / DCM. **^1^H NMR** (400 MHz, CDCl_3_): δ_H_ 7.14 (d, *J* = 2.4 Hz, 2H), 6.62 (t, *J* = 2.4 Hz, 1H), 4.54 (t, *J* = 6.8 Hz, 2H), 4.30 (m, 2H), 4.15-4.03 (m, 4H), 3.95 (t, *J* = 6.4 Hz, 2H), 3.77-3.47(m, 30H), 3.03 (t, *J* = 6.4 Hz, 2H), 2.93 (t, *J* = 6.8 Hz, 2H), 2.20-2.02 (m, 2H), 1.79-1.72 (m, 4H), 1.51-1.18(m, 46H), 0.86 (t, *J* = 6.8 Hz ,6H).

**Synthesis of compound 13**

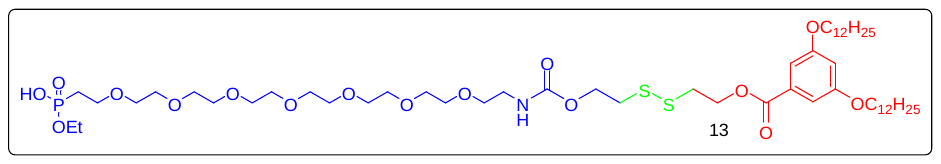

Mol. formula: C_54_H_100_NO_16_PS_2_

Mol. Weight: 1114.47 g/mol

Physical appearance: White solid

Yield: Not determined

The **compound** **13** was prepared from compound **12** (0.070 g, 0.06 mmol) and LiBr (0.050 g, 0.6 mmol). The obtained get crude product, which was used without purification.

**Synthesis of compound 14**

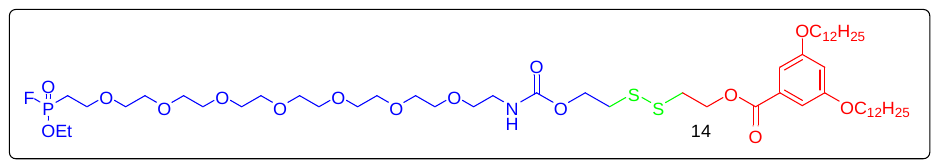

Mol. formula: C_54_H_99_FNO_15_PS_2_

Mol. Weight: 1116.47 g/mol

Physical appearance: White solid

Yield: Not determined

The compound **14** was prepared from compound **13** (0.04 g, 0.03 mmol), DAST (0.024 g, 0.14 mmol. The obtained get crude product, which was used without purification. **^19^F NMR** (400 MHz, CDCl_3_): ^δ^F -59.91, -62.74.

**Synthesis of pH-responsive** **AABP and its intermediates**

**Synthesis of compound 17**

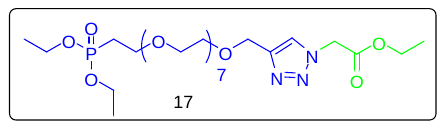

Alkyne **15** (0.18g, 1eq) and Azide **16** (0.5g, 1eq) were dissolved in degassed THF and stirred until clear solution was obtained, then degassed water was added and stirred vigorously for 10 more minutes. Freshly prepared 1M sodium ascorbate (0.028g, 0.05eq) and 1M CuSO_4_ (0.011g, 0.1eq) were added to the reaction mixture at least thrice in an intervals of 45minutes and allowed to react for 16hours at RT. Upon completion, reaction mixture was extracted in DCM and combined organic layer was dried over Na_2_SO_4_ and concentrated under vacuum to get crude product which was purified using normal phase chromatography using MeOH / DCM solvent system. Yield (0.6g, 87%). *R_f_*=0.40 in 5% MeOH / DCM. **^1^H NMR** (400MHz, CDCl_3_): *δ*_H_ 4.16 (d, *J*=2.4Hz, 2H), 3.73-3.62 (m, 15H), 3.22 (t, *J*=7.8Hz, 2H), 2.41 (t, *J*=2.4Hz, 1H). **^13^C NMR (**100MHz, CDCl_3_): *δ*_C_ 79.46, 77.16, 74.61, 71.82, 70.51, 70.29, 70.08, 68.98, 58.30, 3.17.

**Synthesis of compound 18**

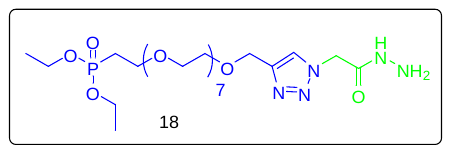

In an oven dried RDF, above ester **17** (100 mg, 1eq) and hydrazine (1mL, excess) were dissolved in absolute ethanol with stirring and refluxed for 12hours. Upon completion of reaction, excess of ethanol and hydrazine was removed under vacuum. To the obtained residue water was added and washed with DCM. Water layer was evaporated under vacuum to get crude product which was purified by column chromatography. Yield (40 mg, 40%). **^1^H NMR** (400MHz, CDCl_3_): *δ*_H_ 7.80 (d, *J*=8Hz, 2H), 7.34 (d, *J*=8Hz, 2H), 4.20-4.14 (m, 5H), 3.70-3.58 (m, 17H), 2.44-2.41 (m, 4H), 1.68 (s, 3H). **^13^C NMR (**100MHz, CDCl_3_): *δ*_C_ 144.89, 129.90, 128.07, 79.71, 74.62, 70.81, 70.65, 70.60, 70.46, 69.33, 69.18, 68.75, 58.47, 21.73.

**Synthesis of compound 20**

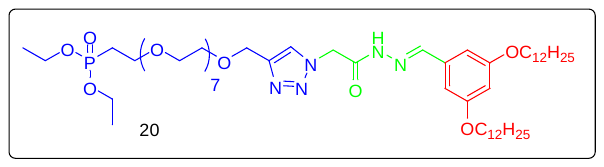

Ethanol was added to the mixture of hydrazine **18** (0.16 g, 1eq) and aldehyde **19** (0.36 g, 2eq) in an oven dried RBF. Then two drops of acetic acid was added and resultant mixture was refluxed for 12hours. Upon completion of reaction, ethanol was removed under vacuum and the obtained residue was loaded and purified using column chromatography using silica gel column chromatography using MeOH / DCM as eluent. Yield (0.115g, 62%). **^1^H NMR** (400MHz, CDCl_3_): *δ*_H_ 7.89 (s, 1H), 7.83 (s, 1H), 6.77 (d, *J*=2.8Hz, 2H), 6.51 (t, *J*=2.4Hz, 1H), 5.65 (s, 2H), 4.75 (s, 2H), 4.15-4.05 (m, 4H), 3.96 (t, *J*=6.4, 4H), 3.90-3.70 (m, 14H), 2.22-2.26 (m, 2H), 1.82-1.73 (m, 4H), 1.73-1.51 (m, 44H), 0.88 (t, H=7.4Hz, 6H).

**Synthesis of compound 21**

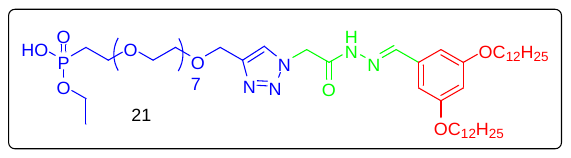

In an oven dried RBF, a mixture of above imine **20** (0.30 g, 1eq) and sodium azide (0.44 g, 10eq) were taken and dissolved in DMF with stirring. The mixture was then heated to 95^⁰^C and allowed to react for 18 hours. Upon completion of reaction, DMF was evaporated under vacuum. Water was added to the residue and extracted thrice with DCM. Upon concentration under vacuum, crude product was obtained which was used without further purification. **^1^H NMR** (400MHz, CDCl_3_): *δ*_H_ 7.87 (s, 1H), 7.81 (s, 1H), 6.78 (d, *J*=2.8Hz, 2H), 6.51 (t, *J*=2.4Hz, 1H), 5.61 (s, 2H), 4.75 (s, 2H), 4.15-4.05 (m, 2H), 3.96 (t, *J*=6.4, 4H), 3.90-3.70 (m, 14H), 2.22-2.26 (m, 2H), 1.82-1.73 (m, 4H), 1.73-1.51 (m, 44H), 0.88 (t, H=7.4Hz, 6H).

**Synthesis of compound 22**

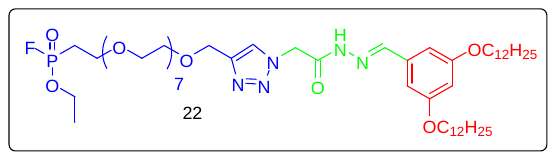

To the stirring solution of monophosphonate ester **21** (25 mg, 1eq) in DCM at -78^⁰^C, DAST (16 mg, 4eq) was added dropwise at RT and allowed to react for 15minutes. Excess of DAST and DCM were evaporated under vacuum. To the obtained residue, was then extracted thrice with DCM. Combined organic layer was dried over Na_2_SO_4_ and concentrated under vacuum to get crude product. **^19^F NMR** (400MHz, CDCl_3_): *δ*_F_ -59.91, -62.74
